# FAK drives resistance to therapy in HPV-negative head and neck cancer in a p53-dependent manner

**DOI:** 10.1101/2022.06.03.494547

**Authors:** Phillip M. Pifer, Liangpeng Yang, Manish Kumar, Tongxin Xie, Mitchell Frederick, Andrew Hefner, Beth Beadle, David Molkentine, Jessica Molkentine, Annika Dhawan, Mohamed Abdelhakiem, Abdullah A. Osman, Jeffrey N. Myers, Curtis R. Pickering, Vlad C. Sandulache, John Heymach, Heath D. Skinner

**Affiliations:** Department of Radiation Oncology, University of Pittsburgh, UPMC Hillman Cancer Center, Pittsburgh, USA; Department of Experimental Radiation Oncology, University of Texas, MD Anderson Cancer Center, Houston, USA; Department of Biochemistry, All India Institute of Medical Sciences (AIIMS), Bilaspur, Himachal Pradesh, India; Department of Head and Neck Surgery, University of Texas, MD Anderson Cancer Center, Houston, USA; Department of Otolaryngology-Head & Neck Surgery, Baylor College of Medicine, Houston, USA; Department of Radiation Oncology, Stanford University, Stanford, USA; Department of Thoracic and Head and Neck Medical Oncology, University of Texas, MD Anderson Cancer Center, Houston, USA

**Keywords:** human papillomavirus negative, head and neck squamous cell carcinoma, head and neck cancer, focal adhesion kinase, p53

## Abstract

Radiation and platinum-based chemotherapy form the backbone of therapy in HPV-negative head and neck squamous cell carcinoma (HNSCC). We have correlated focal adhesion kinase (FAK/*PTK2*) expression with radioresistance and worse outcome in these patients. However, the importance of FAK in driving radioresistance and its effects on chemoresistance in these patients remain unclear. We performed an *in vivo* shRNA screen using targetable libraries to address these questions and identified FAK as an excellent target for both radio- and chemosensitization. Because *TP53* is mutated in over 80% of HPV-negative HNSCC, we hypothesized that mutant *TP53* may facilitate FAK-mediated therapy resistance. FAK inhibitor increased sensitivity to radiation, increased DNA damage and repressed homologous recombination and non-homologous end joining repair in mutant, but not wild-type, *TP53* HPV-negative HNSCC cell lines. Mutant *TP53* cisplatin-resistant cell line had increased FAK phosphorylation compared to wild-type, and FAK inhibition partially reversed cisplatin resistance. To validate these findings, we utilized a HNSCC cohort to show that FAK copy number and gene expression were associated with worse disease-free survival in mutant *TP53*, but not wild-type *TP53*, HPV-negative HNSCC tumors. Thus, FAK may represent a targetable therapeutic sensitizer linked to a known genomic marker of resistance.

## Introduction

Globally, there are greater than 800,000 new cases of head and neck squamous cell carcinoma (HNSCC) a year (1), and curative treatment is associated with profound and life-long toxicity. The only current clinically useful biomarker of outcome in HNSCC is human papillomavirus (HPV), which is associated with better responses to therapy and survival (2). However, patients with HNSCC unrelated to HPV (HPV-negative) continue to have poor clinical outcomes with 3-year progression-free survival between 40%-50% (2). Two of the most critical forms of therapy for this disease are radiation and cisplatin; however, there are no current means by which to personalize therapy using these agents.

One potential avenue for improving response to radiation and chemotherapy is via targeting FAK, a protein frequently overexpressed in HPV-negative HNSCC (3). FAK is a nonreceptor tyrosine kinase involved in cell-cell adhesion (4–6), encoded by gene *PTK2*. At focal adhesions, FAK dimerization results in autophosphorylation at Y397, making FAK catalytically active (5) and leading to phosphorylation of downstream proteins. Our group has previously identified FAK as a driver of radioresistance in HNSCC (3), and FAK has also been implicated in radioresistance, chemoresistance, and immunotherapy resistance in other disease sites (7–10). FAK can be targeted pharmacologically by either FAK and/or dual FAK/PYK2 inhibitors targeting the Y397 site (6). Defactinib is a dual FAK/PYK2 inhibitor currently in clinical trials (6). However, despite this potential importance, the role of FAK inhibition as a therapeutic strategy in HNSCC is unknown. Furthermore, FAK inhibition has not yet been linked to specific genomic drivers of response.

Any potential therapeutic strategy in HPV-negative HNSCC must take into account the tumor suppressor *TP53*, which is the most commonly mutated gene in HPV-negative HNSCC, as well as in many other solid tumors (11, 12). This gene has been associated with negative outcomes in multiple clinical studies of HPV-negative HNSCC, particularly when its predicted function is considered (13–15). FAK and p53 interact at multiple levels. In contrast to its well-known canonical function at focal adhesions, FAK interacts with and regulates p53 in the nucleus where FAK acts as a scaffolding protein and leads to ubiquitation and degradation of p53 (16, 17). Conversely, p53 directly represses FAK transcription (18). Thus, the relevance of FAK in HNSCC may be mediated by the presence or absence of functional p53.

To both examine novel therapeutic sensitizers in HPV-negative HNSCC, as well as assess the impact of *TP53* on therapeutic response, we initially performed an *in vivo* shRNA screen in combination with radiation or platinum-based chemotherapy to identify clinically druggable sensitizing targets in mutant *TP53*, HPV-negative HNSCC. We then hypothesized that wild-type and mutant p53 would have differential effects on FAK activity, DNA damage repair, and responsiveness to FAK-mediated therapeutic sensitization. Finally, we examined the relationship between FAK, *TP53* mutation, and clinical outcome.

## Results

### shRNA screens for radiosensitization and chemosensitization targets

We performed *in vivo* shRNA mediated screens in pre-clinical HNSCC models to discover potential novel agents that were radio- and chemosensitizers. The study of all models for radiosensitization have been published previously (19, 20). In this current study, we focus on mutant *TP53*, HPV-negative HNSCC models from this published screen as well as the same models from a separate experiment in which tumors were treated with carboplatin. Specifically, UM-SCC-22a, HN31, and Cal27 were infected with two barcode labeled shRNA libraries, one consisting of druggable targets and the second focused on DNA damage repair (DDR) proteins (full list of targets in (20) supplemental table 2). These cells were then implanted in the murine flank. Mice were treated with appropriate sham, conventional fractionated radiation therapy (XRT) or carboplatin, with the goal of ~20% tumor volume reduction (19, 20). Subsequently, tumors were harvested, and tumor shRNA bar codes were sequenced. We performed redundant siRNA analysis (RSA), as described previously (19–21). We then evaluated RSA p-values in radiation or carboplatin treated tumors compared to reference cells (generated prior to tumor implant); genes with RSA log p≤-1.3 were considered significant. Of the 502 unique screened targets, 92 genes in both the radiation and carboplatin experiments were significant with 72 out of the 92 genes being associated with response to both carboplatin and radiation (Figure 1A-C). We next performed KEGG pathway analysis of targets found to be significant in both radiation and carboplatin treated tumors, which demonstrated enrichment for genes involved in focal adhesion, proteoglycans in cancer, ErbB signaling pathways, non-small cell lung cancer, cell cycle, and pancreatic cancer (Figure 1D) (22). Finally, we compared targets in untreated tumors versus those treated with either radiation (Figure 1E) or carboplatin (Figure 1F). In this analysis, FAK was preferentially associated with sensitization to both radiation (mean control −1.835 versus mean RT −2.593) and carboplatin (mean carboplatin −2.798).

**Figure 1.**
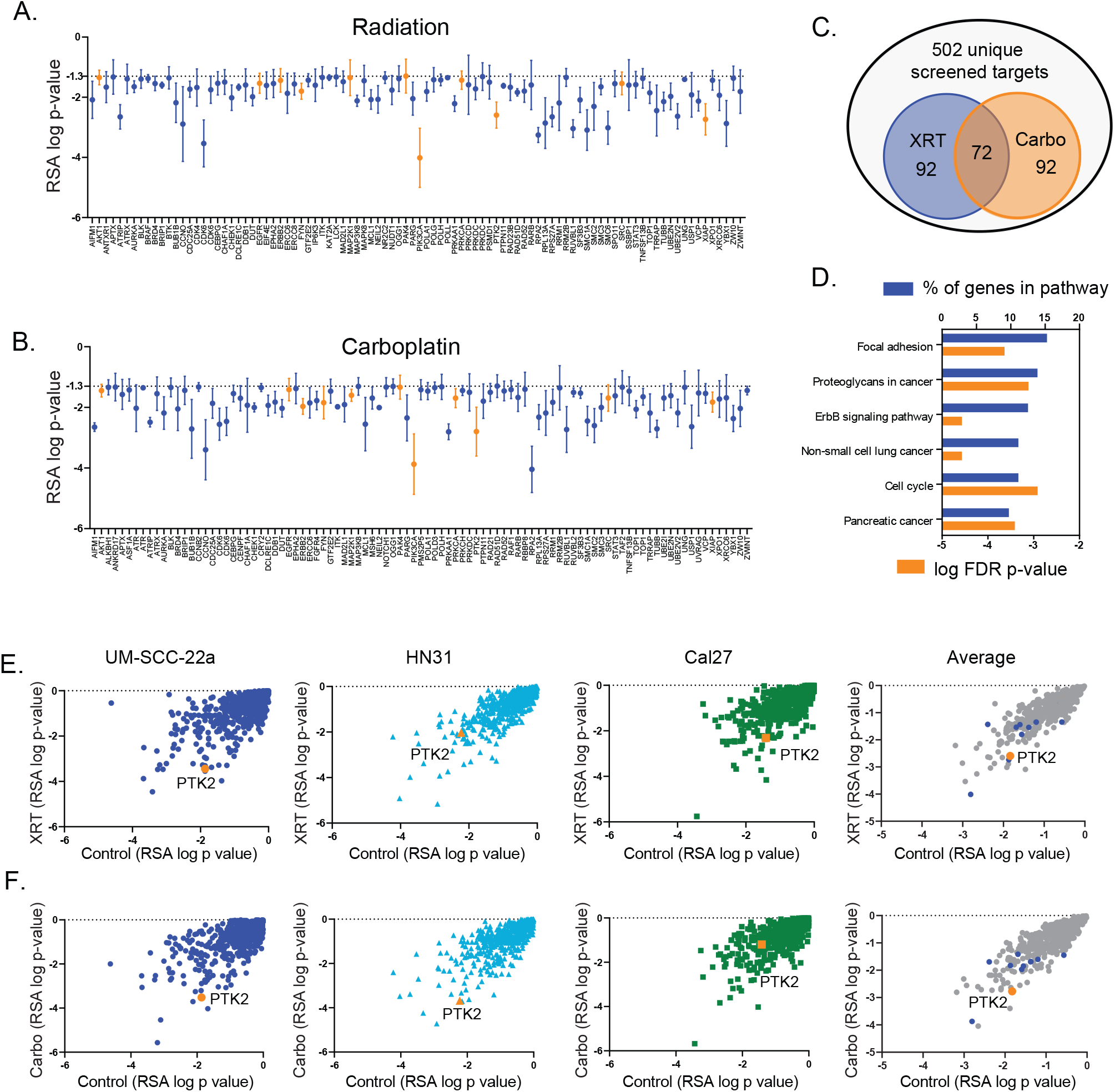
In vivo shRNA screen revealed FAK is a target for radiosensitization and chemosensitization in mutant TP53 HPV-negative HNSCC. A & B) In vivo shRNA screen following treatment with radiation (A) or carboplatin (B) in 3 mutant TP53 HPV-negative tumor models (UM-SCC-22a, HN31, and Cal27). Average RSA-log p-values versus reference for each screened gene shown, with values ≤ −1.3 considered significant. Orange points represent genes in the focal adhesion kinase pathway. C) Venn diagram of genes from the in vivo screen significantly reduced in tumors treated with XRT (n=92), Carboplatin (n=92) or both (n=72). D) Pathway analysis of genes associated with response to both radiation and carboplatin in the in vivo screen (FDR = false discovery rate). E & F) In vivo shRNA screen identified FAK (PTK2) as a radiosensitizing and anti-tumor target in terms of both magnitude of effects (≤ −1.3 log p-value) (E) and as a chemosensitizing and anti-tumor target in terms of magnitude of effects (≤ −1.3 log p-value). All genes shown are statistically significant compared to FDR.

### Radiosensitization by FAK inhibition in a p53-dependent manner

To validate our *in vivo* shRNA screen results for radiosensitization, we performed a murine xenograft experiment using a FAK shRNA knockdown tumor cell line. The combination of FAK inhibition and radiation resulted in a significant decrease in tumor volume compared to FAK inhibition (p < 0.01) or radiotherapy alone (p = 0.01) (Figure 2A). Average area under the curve for HN31 shRNA *in vivo* experiments demonstrated decreasing tumor volume (Control, Control+XRT, shFAK, and shFAK+XRT * p =0.0135) (Figure. 2B).

**Figure 2.**
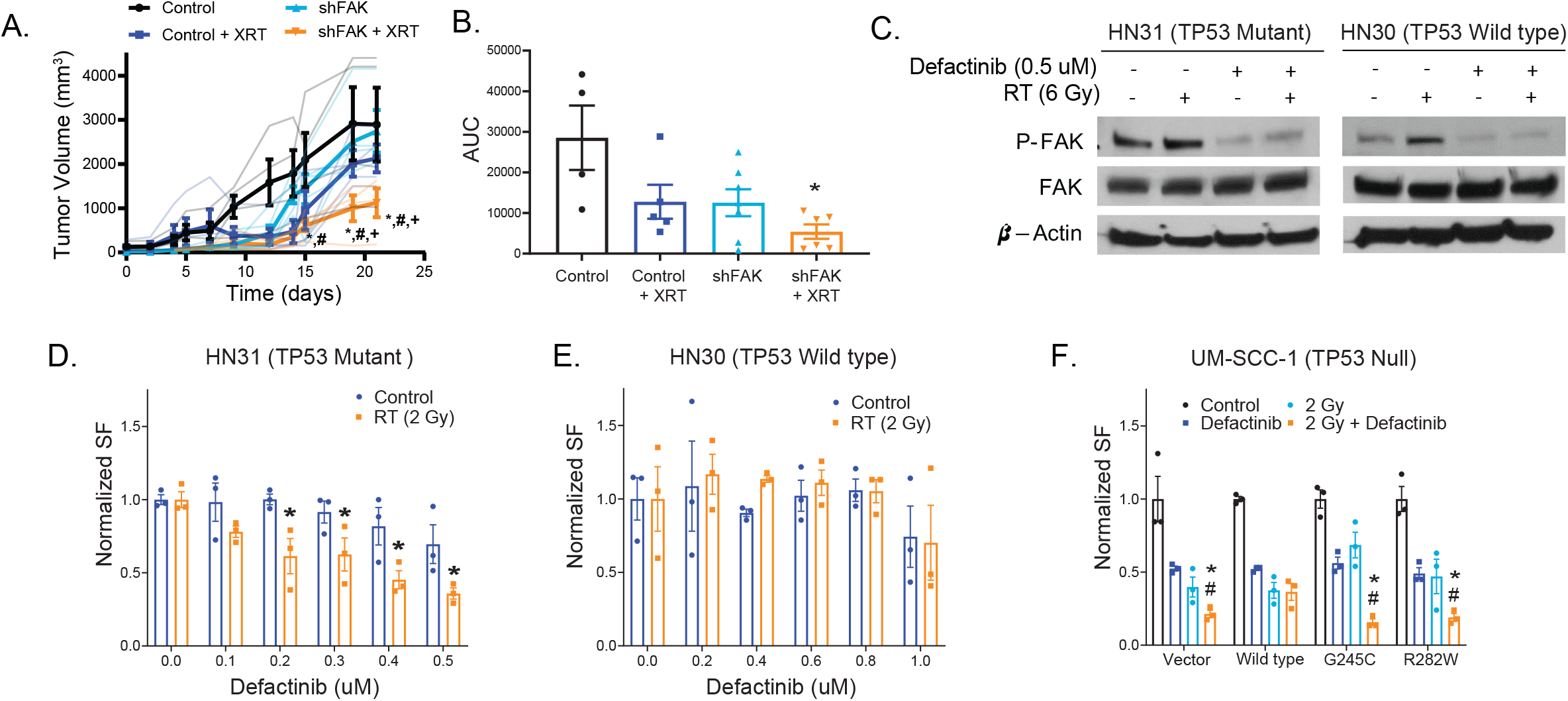
FAK inhibition leads to radiosensitization in mutant, but not wild-type TP53, HPV-negative HNSCC. A) Tumor growth curves for mutant TP53 HPV-negative HN31 xenograft model expressing either control or FAK shRNA in the absence or presence of radiation at 2 Gy for 4 days (Control vs shFAK+RT *- p<0.01, shFAK vs shFAK+RT #, p< 0.01, Control+RT vs shFAK+RT +, p < 0.01). B) Average area under the curve for HN31 shRNA in vivo experiment from (A) (*– p <0.05). C) Immunoblot of mutant TP53 HPV-negative HN31 cells or wild-type TP53 HPV-negative HN30 with pretreatment of 0.5 μM defactinib for 24 hours, irradiation with 6 Gy, and collected at 1 hour time point. D & E) mutant TP53 HN31 (D) and wild-type TP53 HN30 (E) cell lines were treated with defactinib and RT at 2 Gy. Clonogenic survival was determined and normalized to the vehicle treated cells (control vs RT * p <0.05). F) A similar clonogenic survival experiment performed in TP53 null UMSCC-1 cells expressing empty vector, wild-type TP53, or two missense TP53 (G245C and R282W) mutations (Defactinib vs 2 Gy + Defactinib, * - p<0.05; 2 Gy vs 2 Gy + Defactinib #- p<0.05). Individual points within experiments were compared using ANOVA with correction for multiple comparisons. All p-values are two-sided.

Because our *in vivo* shRNA screen was designed to leverage currently available clinically targetable proteins, we next focused on defactinib, a pharmacological ATP-competitive FAK inhibitor. In HPV-negative HNSCC tumor cell lines (HN31 and HN30), defactinib resulted in decreased phospho-FAK, which is a surrogate for FAK activity (Figure 2C). Because the vast majority of HPV-negative HNSCC contain mutant *TP53*, we investigated if FAK inhibition was dependent on *TP53* mutational status. Our model system used isogenic HNSCC cell lines from a primary tumor (wild-type *TP53* HN30) and lymph node metastasis (mutant *TP53* HN31) (23). In clonogenic survival assay, the mutant *TP53* cell line (HN31) was radiosensitized to radiation by defactinib, whereas the wild-type *TP53* HN30 cell line survival fraction remained unchanged with increasing doses of defactinib (Figure 2D&E). Similarly, when we expressed wild-type and mutant (G245C and R282W) *TP53* in a *TP53* null cell line (UMSCC1), the combination of FAK inhibition and radiation did not have an additive effect in wild-type *TP53* cells but resulted in radiosensitization in the null and mutant *TP53* cell lines compared to both radiation and defactinib alone at comparable levels (Figure 2F).

### Chemosensitization by FAK inhibition in a p53-dependent manner

To validate our *in vivo* shRNA screen results for chemosensitization, we inhibited FAK via shRNA or defactinib in a clonogenic survival assay. FAK inhibition resulted in chemosensitization in the presence of carboplatin (Figure 3A&B). Immunoblot analysis in HNSCC cell lines demonstrated that FAK inhibition resulted in decreased phospho-FAK in the presence of carboplatin (Figure 3C). To examine the role of platinum-based chemoresistance, *TP53* mutational status, and FAK inhibition, we initially performed MTT assays combining carboplatin and defactinib in both wild-type *TP53* HN30 and mutant *TP53* HN31 cells (Suppl figure 1). Generally, modest sensitization to carboplatin by defactinib was observed in HN31 cells, with no sensitization observed in HN30 cells. We additionally evaluated HN31 and HN30 cell lines that were rendered cisplatin-resistant (HN31-CR and HN30-CR) due to serial passage in cisplatin-containing media (24). The HN30-CR cell line still contains wild-type *TP53*.Phospho-FAK expression was higher in the HN31-CR cells compared to the parental cell line; however, this was not observed in the HN30-CR line (Figure 3D). Moreover, treatment with defactinib significantly sensitized the HN31-CR cells to cisplatin, while FAK inhibition had no effect on HN30-CR cisplatin response in MTT assays (Figure 3E, F).

**Figure 3.**
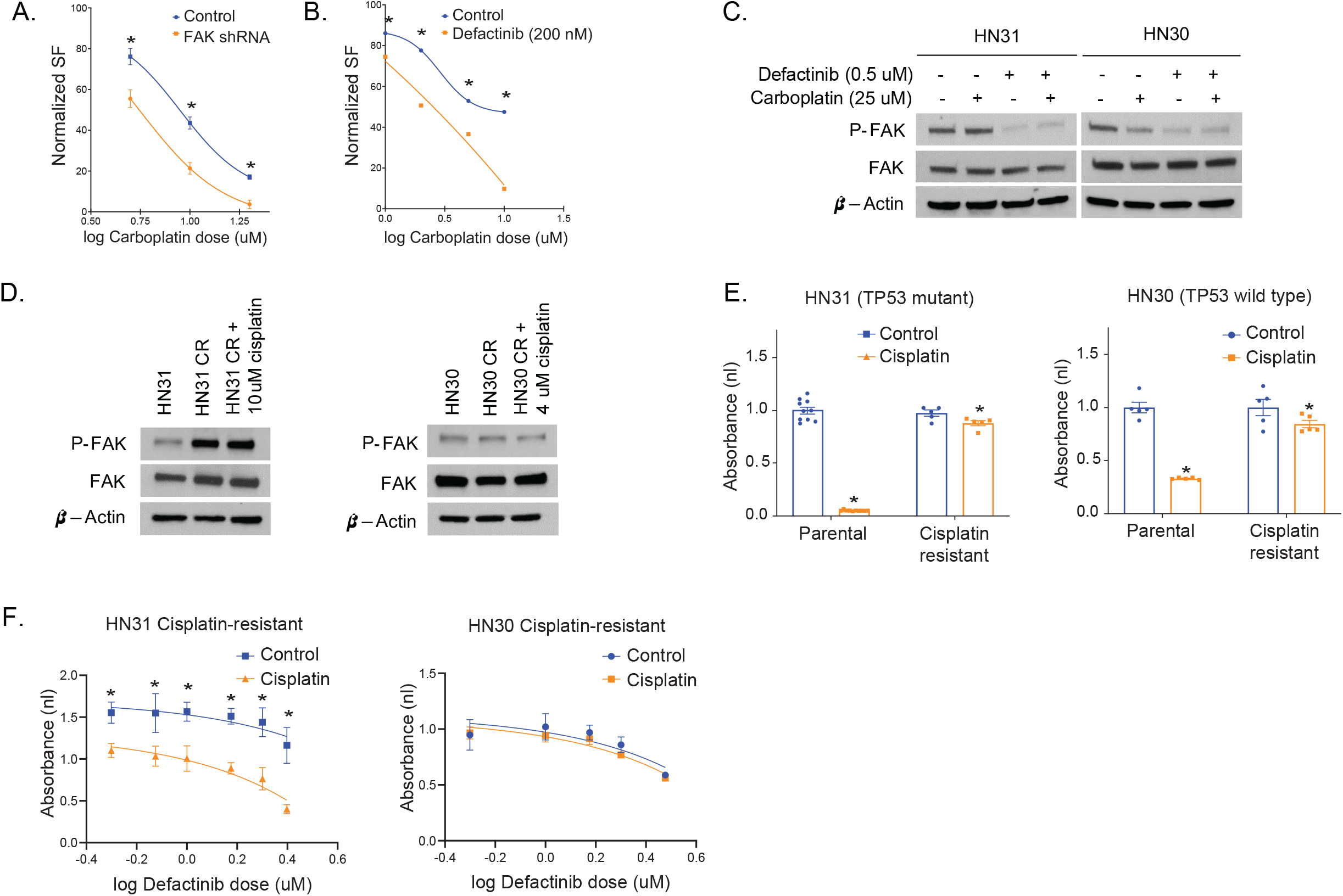
FAK inhibition leads to chemosensitization in mutant TP53 HPV-negative HNSCC. A & B) Mutant TP53 HPV-negative HN31 cell lines were treated with carboplatin combined with either FAK shRNA (A) or defactinib (B) and carboplatin. Clonogenic survival was determined and normalized to the vehicle treated cells. C) Immunoblot of mutant TP53 HN31 cells or wild-type TP53 HN30 cells pretreated with defactinib for 24 hours, followed by the addition of carboplatin for 24 hours. D) Immunoblot of mutant TP53 HN31 and wild-type TP53 HN30 parental and cisplatin resistant (CR) cell lines treated without or with cisplatin. E) MTT assay of mutant TP53 HN31 and wild-type TP53 HN30 parental and cisplatin resistant (CR) cell lines treated with indicated doses of cisplatin and defactinib normalized to 0uM Cisplatin and 0 uM defactinib. F) MTT assay of mutant TP53 HN31 and wild-type TP53 HN30 cisplatin resistant (CR) cell lines treated with indicated doses of cisplatin and defactinib normalized to cisplatin dose. ANOVA with correction for multiple comparisons was used. * Denotes two-sided p<0.01.

### FAK Inhibition and DNA Damage Response

We next focused on the relationship between *TP53* status and FAK inhibition as it relates to oxidative stress and DNA damage, as these processes generally underlie the cellular response to both platinum-based chemotherapy and radiation. Using UMSCC1 cells expressing wild-type or mutant (G245D) *TP53*, cell lines were treated with defactinib, radiation, or the combination. Mutant *TP53* cell line demonstrated increased basal phospho-FAK levels compared to wild-type *TP53* (Figure 4A).

**Figure 4.**
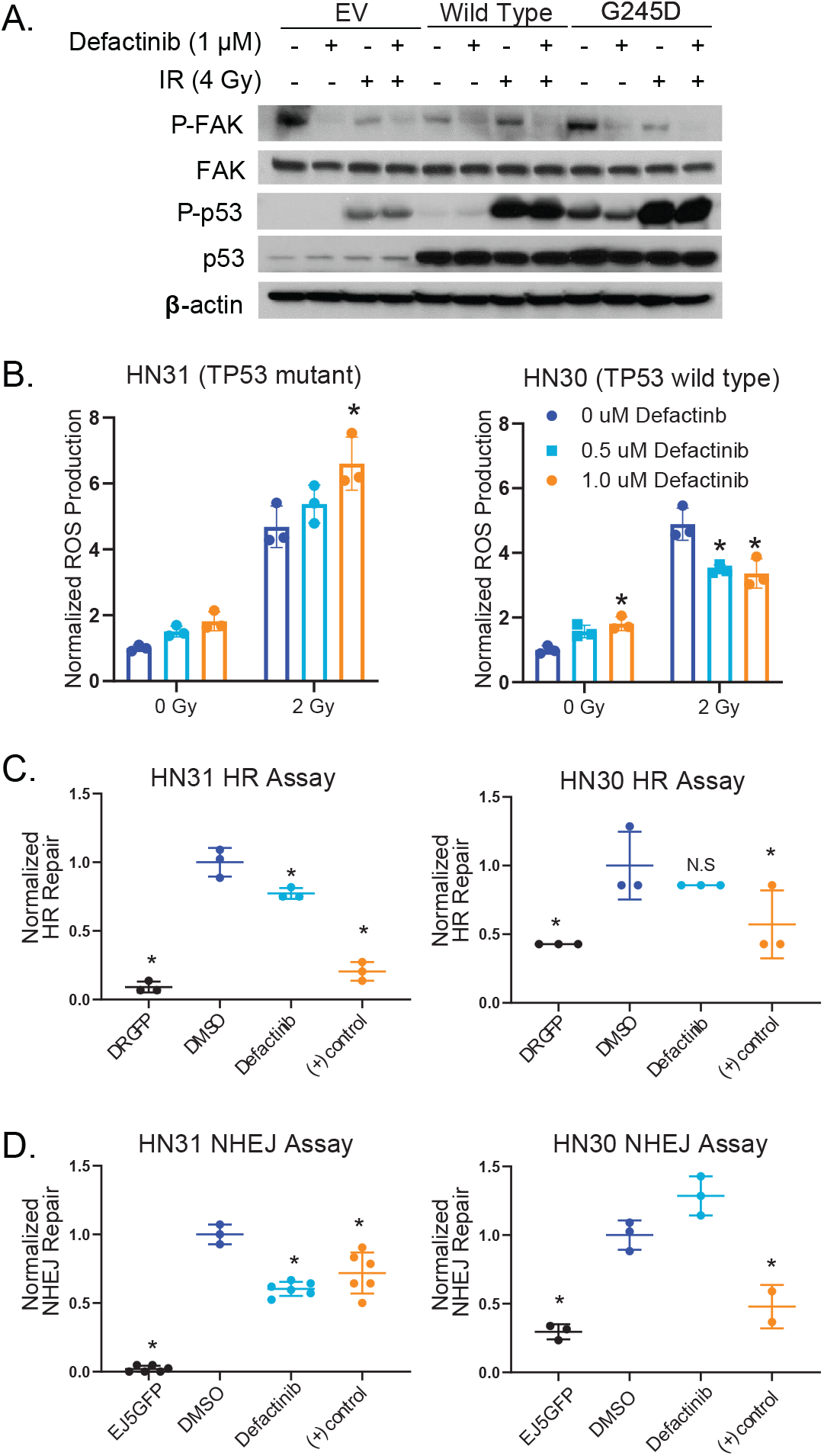
FAK phosphorylation, reactive oxygen species (ROS) and DNA damage repair are differentially affected by TP53 status in HPV-negative HNSCC. A) Immunoblot of UM-SCC-1 cells forced to express wild-type or mutant (G245D) p53 treated with 1.0 μM Defactinib and ionizing radiation (4 Gy) (B) ROS production in mutant TP53 HN31 and wild-type TP53 HN30 cells following treatment with defactinib and ionizing radiation (2 Gy) measured via DCF-ROS flow cytometry assay. C-& D) I-Scel assay for HR (C) and NHEJ (D) in mutant TP53 HN31 and wild-type TP53 HN30 cells following treatment of defactinib. HR= homologous recombination and NHEJ= nonhomologous end joining. ANOVA with correction for multiple comparisons was used. * – denotes two-sided p<0.01.

When we examined the combination of radiation and defactinib, we found significantly increased ROS (as measured by DCFDA assay) in mutant *TP53* HN31 cells (Figure 4B), with similar results seen in a separate mutant *TP53* cell line (FADU, not shown). Conversely, we found a repression of ROS in the wild-type *TP53* HN30 cells using the same combination treatment (Figure 4B).

We next evaluated the effect of FAK inhibition on homologous recombination (HR) and nonhomologous end joining (NHEJ) DNA damage repair via a I-Scel-based reporter assay (20, 25). Pharmacological FAK inhibition in mutant *TP53* HN31 cells resulted in significant decreases in both NHEJ (p = 0.02) and HR (p = 0.04), but FAK inhibition in wild-type *TP53* HN30 cells resulted in no significant effects on NHEJ (p = 0.13) and HR (p = 0.42) (Figure 4C&D).

### Clinical outcomes of patients with mutant *TP53* HPV-negative HNSCC and FAK expression

To investigate the clinical relevance of FAK in HPV-negative HNSCC, we evaluated pre-treatment tumor specimens from patients treated with postoperative radiation therapy (3, 26). Patients were stratified by *TP53* mutational status (wild-type versus mutant) and either *PTK2*/FAK copy number (amplification, gain, or no change/deletion) or *PTK2*/FAK mRNA expression (high vs. low). FAK amplification or gain was significantly associated with worse disease-free survival (DFS) in the mutated *TP53* patient cohort (p =0.02), but not in the wild-type *TP53* cohort (p=0.20). Similarly, stratification of patients by high or low FAK expression and *TP53* mutational status demonstrated worse disease-free survival in the patients with high FAK expression and mutated *TP53* compared to other groups (p=0.013) (Figure 5).

**Figure 5.**
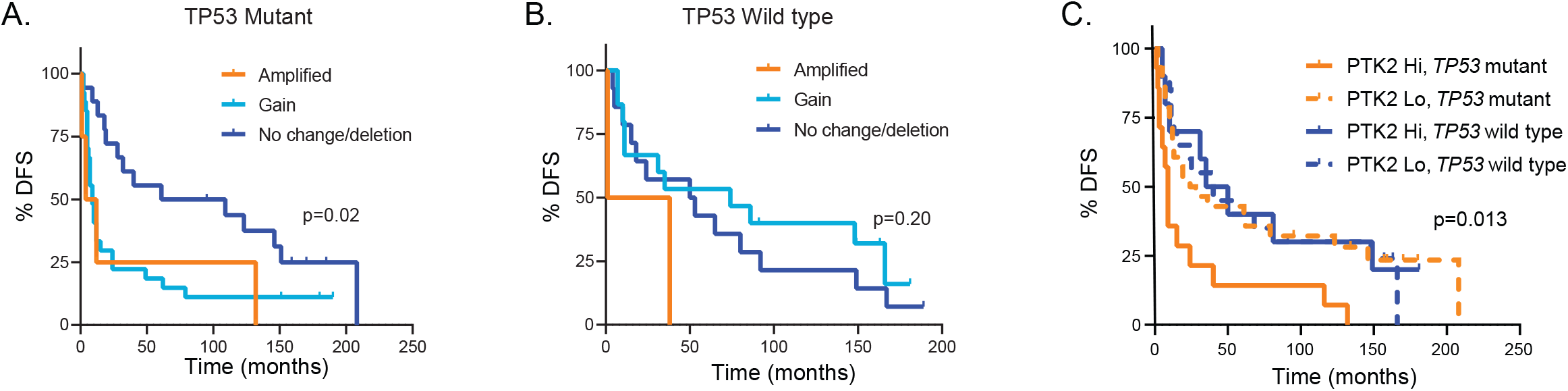
FAK is associated with worse disease-free survival (DFS) following radiation in mutant TP53 HPV-negative HNSCC. A & B) HPV-negative tumors treated with surgery and post-operative radiation (n=94) for the institutional cohort were separated by into either TP53 mutant (A, n=49) or wild-type (B, n=31). Each cohort was then further stratified by PTK2 copy number as performed previously. C) Patients were stratified by TP53 status (mutant versus wild-type) and PTK2 mRNA expression (upper tertile vs. others) as described in the Methods. Statistical analysis via Kaplan-Meier with log rank analysis. Two-sided p-values are shown in the figure.

## Discussion

Currently, the only clinically used biomarker in HNSCC is HPV, the presence of which is associated with improved outcome compared to its HPV-negative counterpart (2). Thus, biomarkers in HPV-negative HNSCC are desperately needed to improve prognostication and identify patients appropriate for treatment intensification and additional systemic therapies. *TP53* mutational status is a natural candidate for this role since it is the most commonly mutated gene in HPV-negative HNSCC and has been previously associated with worse clinical outcome (13, 14, 27). Unfortunately, the use of *TP53* mutation as a biomarker in HNSCC is complicated by the fact that *TP53* mutations have differential effects ranging from no effect to a loss or even gain of function (28). This phenomenon has led to the use of empirical methods of classifying *TP53* to define clinical outcome, without a satisfactorily validated biomarker (13, 14, 27). However, building on our previous work linking FAK to radioresistance in HNSCC, we found that combining FAK with a simple binary classification (wild-type vs mutant) of *TP53*, we improved the ability to identify patients resistant to current therapy for HNSCC.

Our observation linking FAK and *TP53* status is particularly impactful in that we identified FAK as a highly significant target for sensitization to both radiation and platinum-based chemotherapy in mutant *TP53* HPV-negative HNSCC using an *in vivo* shRNA screen. This screen utilized a medium throughput evaluation of targetable genes in an *in vivo* setting that allows for prioritization of targets based on their relative effects on therapeutic response, with FAK – and its associated signaling cascade – generally ranking high for both radio- and chemosensitization in most tumor models tested. By design, our screen included genes that are targetable with currently available agents. While FAK inhibition is a suboptimal treatment in the monotherapy setting, it appears well-tolerated as a single agent in early phase clinical trials, and combinatorial trials with other anti-neoplastic agents are ongoing (6, 29–32). Thus, our data link tumors most resistant to therapy with a clinically viable strategy, namely FAK inhibition, that can sensitize to both radiation and chemotherapy.

On its face, the idea that FAK inhibition might sensitize broadly to therapies that work through DNA-damage may be non-intuitive. FAK is a crucial signaling step between β1-integrin and growth factor signaling and the intracellular signaling cascade that leads to reorganization of the actin skeleton, lamellipodia formation, and proliferation (5, 33). Thus, FAK has been approached as a therapeutic target largely on the basis of reducing cellular motility and metastasis. However, previous work has implicated β1-integrin signaling in resistance to radiation in HNSCC, in a manner potentially dependent upon cell-cell interaction (34–36). Our own work, and that of others, also directly implicates increased intracellular ROS and repression of DNA-damage repair following FAK inhibition with therapeutic sensitization (3, 9, 35). The exact mechanism of this phenomenon is likely multifactorial but remains to be completely elucidated.

Separately we found that FAK-mediated therapeutic sensitization was dependent upon the presence of mutated *TP53*. Wild-type p53 is known to repress the transcription of PTK2 at the level of the promoter in some models, a phenomenon not observed when mutant TP53 was expressed (18). Additionally, FAK can repress the function of p53 via repression of p53 transcription. Therefore, a feedback mechanism exists between these signaling pathways at the transcriptional level (37). However, similar to Boudreau and colleagues, we did not observe a relationship between total FAK levels and *TP53* status but did see repression of FAK phosphorylation in cells expressing wild-type p53 versus mutant p53 isoforms (38). Moreover, in the mutant *TP53* cells rendered resistant to cisplatin, we observed a dramatic increase in FAK phosphorylation, but not total FAK level, compared to the parental line, that was not observed in the wild-type p53 setting. These lines of evidence suggest that the critical effects of FAK modulation on therapeutic response may lie at the post-transcriptional level. Indeed, it has been shown that wild-type p53 can bind FAK directly, which can potentially repress the function of both proteins (17, 39, 40). We have additionally found that at least some forms of mutant p53 can bind to FAK at levels similar to wild-type, however without a commensurate decrease in FAK activity. It is likely that in tumors with mutant *TP53*, high levels of FAK activity are attributable to the absence of functional p53 and makes tumors more sensitive to the effects of FAK inhibition.

### Conclusion

Our *in vivo* shRNA screen using radiation and platinum-based chemotherapy demonstrated FAK inhibition as a potential target for both radiosensitization and chemosensitization. *In vivo* and *in vitro* data suggest that the benefit occurs in mutant *TP53* HPV-negative HNSCC which is supported by clinical outcomes.

## Material and methods

### *In Vivo* shRNA Screen and analysis

The *in vivo* shRNA screen method was described previously (19, 20, 41). For this analysis, three HPV-negative HNSCC cell lines (UM-SCC-22a, HN31, and Cal27) were used. Briefly, lentiviral particles of previously generated druggable proteins and proteins involved in DNA-damage repair shRNA libraries were infected *in vitro* using spinfection into the HPV-negative HNSCC cell lines at a low MOI. Cells were selected with puromycin, and four million cells were subcutaneously injected into the flank of nude mice, and reference cells from the day of injection were collected and frozen. Once the tumor had reached approximately 100 mm^3^, the xenografts were treated with 2 Gy/day of radiation or varying doses of carboplatin, with a goal of achieving ~20% tumor volume reduction. The tumors were subsequently allowed to grow to ~ 500 mm^3^, harvested, and DNA was isolated. Tumor and references cells were then amplified and sequenced on Illumina sequencers (41). Analysis was performed as detailed in Carugo *et al* and Kumar *et al* for detailed analysis please refer to (20, 41). Briefly, shRNA hairpin counts were normalized to counts per million, and log2 fold-change for each shRNA hairpin was calculated compared to reference pellet shRNA hairpin level. A modified version of siRNA activity (RSA) algorithm (42) was used to generate gene-level summary measure per cell line with modifications to ensure that at least two hairpins were used to calculate p values and hairpins that ranked above luciferase controls were excluded. Data was displayed graphically using GraphPad Prism (v8.0).

### Cell Lines

HN30, HN31, and UMSCC-1 cell lines were generously supplied by Dr. Jeffrey Myers via the University of Texas MD Anderson Cancer Center Head and Neck cell line repository. HN30 and HN31 cisplatin-resistant cell lines were generously supplied by Dr. Vlad Sandulache from Baylor College of Medicine. HN31 cell line with control and FAK shRNA generation were previously described (3). Before experiments, cell lines were genotyped and tested for mycoplasma. Cell lines were cultured in DMEM/F-12 50/50 medium supplemented with 10% fetal bovine serum and 1% penicillin/streptomycin and incubated at 37°C and 5% CO_2_ atmosphere. HN30 and HN31 cisplatin-resistant cell lines were cultured in 4 μM and 10 μM of cisplatin, respectively. Defactinib (Cat. #S7654), Cisplatin (Cat. # S1166) and Carboplatin (Cat. # S1215) were purchased from Selleck Chemicals (Houston, TX).

### Clonogenic Survival Assay

Assays were performed as previously described (20). Briefly, single cells were plated in a 6 well plate overnight. The next day cells were pre-treated with DMSO or indicated dose of defactinib for 24-hours for experiments. Cells were then irradiated at indicated dose or treated with carboplatin for 24-hours. Cells were re-fed with media. The cells formed colonies over the next 10-14 days. Colonies were fixed with 0.25% crystal violet/methanol solution, and colonies containing greater than 50 cells each were counted (20). Clonogenic survival curves were generated using GraphPad Prism (v8.0).

### 3-[4,5-dimethylthiazol-2-yl]-2,5 diphenyl tetrazolium bromide (MTT) assay

The MTT assay was performed similar to previous publication with the following modifications (43). Depending on cell line, 1,000-4,000 cells per well in 100 μL of media were plated into 96-well plates. After 24-hour incubation, cells were treated with 100x defactinib, carboplatin, cisplatin and/or appropriate control. After 48 hours, cells were aspirated and re-fed with media. Once control cells were confluent (~48-72 hours after refeeding), 50 μL of 5 mg/mL Thiazolyl Blue Tetrazolium Bromide solution (Sigma, MTT) was added to cells and incubated for 3 hours. Wells were aspirated and 150 μL DMSO was added to each well. 96 well plates were agitated for 30 minutes on shaker. Plates were read using an Epoch Microplate Spectrophotometer (BioTek) at 590 nm and Gen5 v.3.05 software (Gen5, RRID:SCR_017317). Plots were generated using GraphPad Prism (v8).

### Non-homologous end joining and homologous recombination DNA Damage Repair Assay

HN30 and HN31 cell lines with pDRGFP for HR assay (Addgene, Plasmid #26475) or EJ5GFP for NHEJ assay (Addgene, Plasmid 44026) were generated as previously described (20). The HN30 and HN31 cell lines were transfected with 2 μg mCherry (Addgene, Plasmid 41583) as a negative control, or with both 2 μg mCherry and 6 μg pCBASceI (Addgene, Plasmid 26477). The cells were incubated overnight in 1 ml of media containing DMSO for the controls, 500 μM Defactinib, or 50 uM ATMi for HN30 and 10 mM ATMi for HN31 as the positive controls. The media was replaced with 2 ml of media the following day with the same concentrations of drug added for 24 hours. The cells were then trypsinized, centrifuged into a pellet at 1200 rpm for 5 minutes, and washed with PBS before being resuspended in 1 mL of FACS buffer containing PBS, 0.1% BSA (Sigma) and 0.1% NaA (Sigma). Flow cytometry was run using the BD Accuri C6 Plus flow cytometer (BD Biosciences) to detect GFP and RFP, with cells positive for mCherry and GFP being gated as positive (20). The data was plotted with GraphPad Prism.

### Reactive oxygen species (ROS) measurement

Intracellular ROS levels were measured according to previously published protocol using CM-H_2_DCFDA (ThermoFisher Scientific, Cat: D399) (44, 45). Briefly, cells were treated with DMSO control or defactinib at indicated doses for 24-hours. Cells were irradiated at indicated dose. Cell culture media was removed, and cells were stained in serum-free medium containing 5 μM H_2_DCFDA for 30 minutes at 37°C. Positive control (serum-free medium with 5 μM H_2_DCFDA and 100 μM H_2_O_2_) and negative control (no H_2_DCFDA) were used for all experiments. Cells were trypsinized and harvested by centrifugation at 400g at 5 min. Cells were washed once with PBS and centrifuged as above. Cells were suspended in 0.5 ml of PBS and assayed immediately by flow cytometry with fluorescence excitation and emission (492-495 / 517-527 nm).

### *TP53* Expression

*TP53* constructs (wild-type, G245D, R175H) in pBabe retroviral expression vector (pBABEpuro; Addgene) were a generous gift from Drs. Jeffery Myers and Abdullah A. Osman from the University of Texas MD Anderson Cancer Center (15). Retroviral packaging performed in GP2+293T cells by transfecting 4.25 μg of *TP53* constructs and 1.75 μg of CMV-VSG-G with 18 uL of polyethylenimine hydrochloride in DMEM/F12 media with 1% FBS for 6 hours, and then cells were re-fed. *TP53* construct virus-containing media was harvested from GP2-293T cells at 48 hours and passed through 0.45 μm syringe filter. Three mL of *TP53* construct virus-containing media with 5 μL of polybrene was placed on UMSCC1 cells in 6-well plate for 6 hours, and the virus containing media was replaced with growth media. This entire process was repeated after 48 hours. *TP53* construct UMSCC1 cells were then selected with 2 μg/mL puromycin, and *TP53* expression was confirmed via western blot.

### Mouse Xenograft Model

The FAK shRNA *in vivo* study was performed according to all relevant ethical regulations after IACUC approval from the University of Texas MD Anderson Cancer Center similar to previous xenograft models (20). Male athymic nude mice (6 weeks old, Cold Spring Harbor) were injected with HN31 tumor cells with control or FAK shRNA (2 × 10^6^ in 0.1 ml of PBS) subcutaneously in the right dorsal flank of each mouse. Tumor diameters were measured with calipers, and tumor volume was calculated as A × B2 × 0.5 (A= largest diameter, B=shortest diameter). When the tumors volumes reached ~100-150 mm^3^, mice were balanced according to tumor size into sham or ionizing radiation (2 Gy daily x 4 days) and subsequently tracked for three weeks for tumor-growth delay experiments. Tumor volume was measured every 2-3 days (calculated as above) and averaged between groups. Group comparisons were performed using Two-Way ANOVA with post-hoc false discovery rate (FDR). Statistical analysis was performed in GraphPad Prism v8.

### Immunoblot Analysis

Immunoblot analyses were performed as previously described (20). Cell lines were washed with cold PBS and scraped into whole-cell lysis buffer (20 mM HEPES, pH 7.9, 0.4 M NaCl, 0.1 mM EDTA, pH 8.0, 0.1 mM EGTA, pH 7, 1% Igepal, 1X Halt protease-inhibitor cocktail (Thermo Sci), and 1x Halt phosphatase-inhibitor cocktail (Thermo Sci). Protein lysates were vortexed and sonicated at 100 amplitude for 2 minutes using QSonica Q700 sonicator (Newton, CT). Subsequently, protein lysates were centrifuged for 15 mins at 4°C, and supernatant were transferred to microcentrifuge tube. Protein assays were performed using DC Protein Assay kit (BioRad) and was used to load equivalent total protein on 4-15% gradient (SDS)-polyacrylamide gel (Bio-Rad). Protein was transferred for 10 min onto polyvinylidene-difluoride membrane using Transblot Turbo device (Bio-Rad). Membranes were blocked with 5% nonfat powdered milk in TBS (TBS, 0.1 M, pH=7.4) and incubated with primary antibody at 4°C overnight. Primary antibodies used include: phospho-FAK (Tyr397,1:250, #3283), total FAK (1:1000, #3285), phospho-histone H_2_A.X (Ser139, 1:500, #2577S), histone H_2_A.X (1:1000, #2595), phospho-p53 (Ser20, 1:500, #9287), p53 (1C12, 1:1000, #2524), p21Waf1/Cip1 (12D1, 1:1000, #2947), phospho-SAPK/JNK (Thr183/Tyr185, 1:500, #9251), and phospho-CHK2 (Thr68, 1:500, #2661) from Cell Signaling Technology (Danvers, MA); β-actin (C4, 1:10,000, #MAB1501) from MilliporeSigma (Burlington, MA): and p53 (DO-1, 1:1000, #SC-126) from Santa Cruz Biotechnology (Santa Cruz, CA). Membranes were washed three times with TBS + 0.1% Tween and incubated with goat anti-rabbit (1:2000, #NA934V) or anti-mouse (1:2000, #NA931V) secondary antibodies conjugated to horseradish peroxidase (GE Healthcare, Chicago, Illinois) at room temperature for 1 hour. Membranes were washed as above, and signal was generated using ECL2 western blotting substrate (Pierce Biotechnology, Rockford, IL) on Hyblot CL autoradiographic film (Thomas Scientific, Swedesboro, NJ).

### Immunoprecipitation

Immunoprecipitation was performed as previously described (20). Briefly, cell lines were grown in 15-cm dished and lysed in 1ml of Pierce IP lysis buffer with 1X Halt protease-inhibitor cocktail (Thermo Sci), and 1x Halt phosphatase-inhibitor cocktail (Thermo Sci) on ice. Protein lysates were sonicated at 100 amplitude for 2 minutes using QSonica Q700 sonicator (Newton, CT), and lysates were centrifuged for 15 mins at 4°C, and supernatants were transferred to prechilled microcentrifuge tubes. Protein assay was performed using DC Protein Assay kit (BioRad). One mg of total protein of each sample were immunoprecipitated with 50 ul of p53 antibody (D0-1, sc-126, Santa Cruz) and incubated overnight on rotating wheel at 4°C, and 50 ul of 100mg/ml Protein A Sepharose Beads (17-0780-01, GE Healthcare) were added to the samples and incubated for 2 hours on rotating wheel at 4°C. Samples were then washed with IP lysis buffer three time and boiled in 25 ul of 2X SDS-loading buffer for 5 minutes and loaded into 4-15% polyacrylamide gels (BioRad).

### Clinical Data

An institutional cohort of HPV-negative HNSCC treated with surgery and post-operative radiation was examined, with tumors evaluated via Illumina gene expression array and targeted sequencing as described previously (3, 26). A total of 94 patients had tumors with detailed clinical outcomes, gene expression, and *TP53* status available. Patients were stratified by *PTK2*/FAK linear copy number as performed previously (3) and separated into wild-type and mutant *TP53* groups for analysis. *PTK2* mRNA expression for wild-type and mutant *TP53* tumors was assessed, and patients were stratified into either *PTK2* high (upper tertile) or low (remaining tertiles) groups for outcome analysis. Disease-free survival curves were generated by using the method of Kaplan-Meier, with log rank statistics used to determine significance. SPSS (v27) was used for clinical analysis, with figures generated using Graph Pad Prism (v8).

## Statistical Analysis

*In Vivo* shRNA screen statistical analysis was performed as described in section above. Clonogenic survival curves were analyzed using two-way ANOVA with post hoc analysis for multiple comparisons. ROS, HR, and NHEJ assays were analyzed using a paired t-test. Tumor growth curves for HN31 xenograft model were analyzed using matched mixed-effects model, and average area under the curve for HN31 shRNA *in vivo* experiment was analyzed using ordinary one-way ANOVA. For clinical data, disease-specific survival was generated by using the method of Kaplan-Meier with log rank statistics, and p< 0.05 was considered significant unless otherwise specified. Statistical analysis was generated using GraphPad Prism (v8.0).

## Study Approval

Murine experiments were performed according to all relevant ethical regulations after IACUC approval from The University of Texas MD Anderson Cancer Center.

## Author contributions

**Conceptualization:** HS, PMP, LY, MK, LP, TX, MF, CP, JM

**Methodology:** HS, PMP, LY, TX, CP, MF, BB, MA

**Validation:** MK, PMP, AH, LY

**Formal analysis:** HS, PMP

**Investigation:** PMP, MK, LY, TX, JM, DM, AH, AD

**Resources:** HS, JM, AAO, JH, VS

**Data Curation:** HS, PMP, LY, TX

**Writing - Original Draft:** PMP, HS

**Writing - Review & Editing:** HS, PP, LY, MK, LP, TX, MF, CP, JM, AAO, JH, VS, BB, MA

**Visualization:** HS, PMP

**Supervision:** HS, CP, JM, JH, MF

**Project administration:** HS, JM, MF, JH

**Funding acquisition:** HS, CP, JM

## Acknowledgments

This study was supported by: 1) the National Cancer Institute R01CA168485-08 (HS) and T32 CA060397-26 (PMP) (2) the National Institute for Dental and Craniofacial Research R01 DE028105 (HS), R01DE028061 (HS and CP), and U01DE025181 3) The Cancer Prevention Institute of Texas RP150293 (HS and CP) and 4) Veterans Administration Clinical Science Research and Development Division Career Development Award 1IK2CX001953 (VS).

---

**Supplemental Figure 1.**
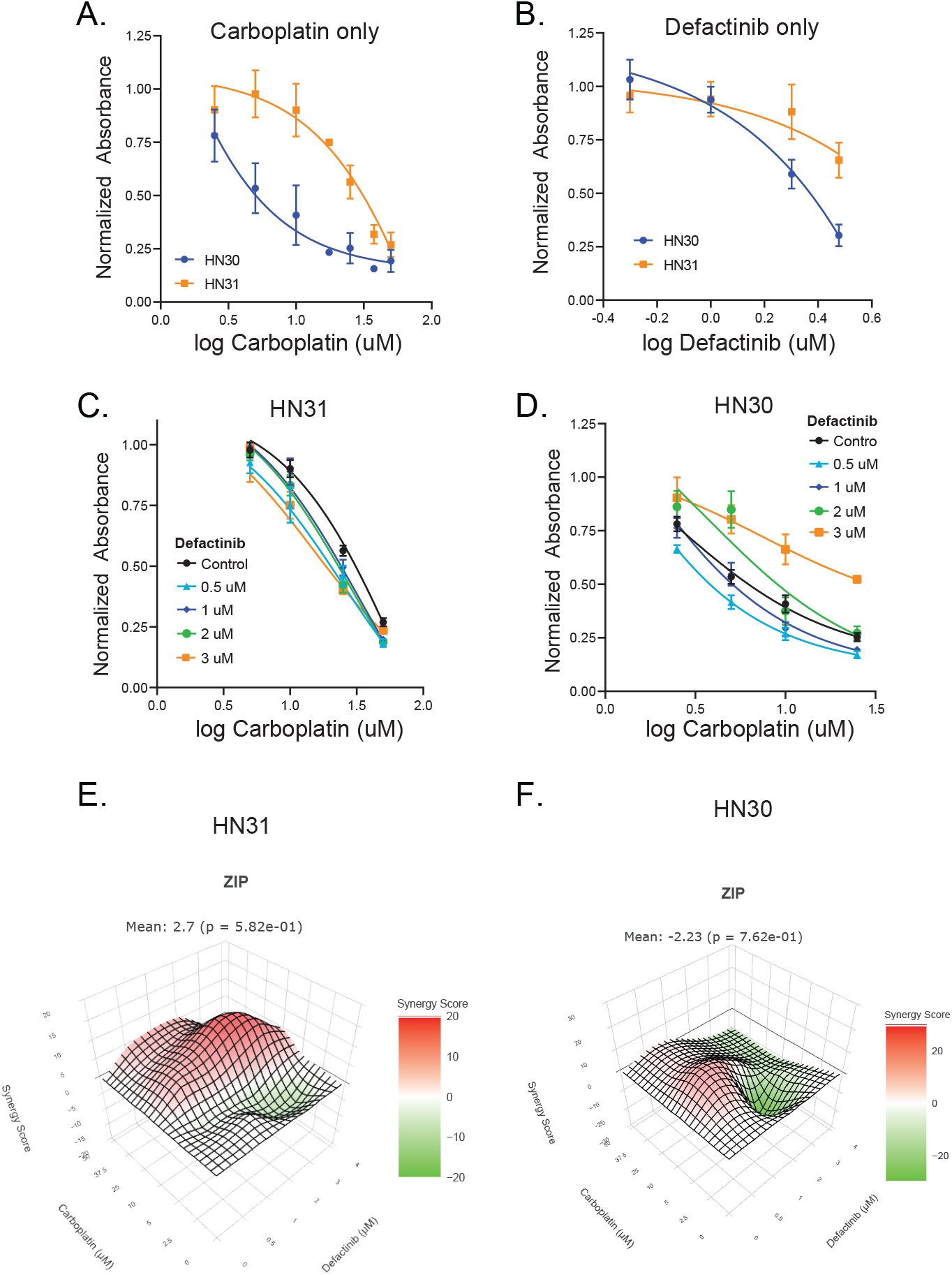
FAK inhibition in HN30 and HN30 cell lines treated with carboplatin. A & B) MTT assay in HN31 and HN30 cells treated with (A) carboplatin or (B) defactinib. C & D) MTT assay in HN31 (C) and HN30 (D) cells treated with the combination of defactinib and carboplatin at the indicated doses. E & F) ZIP Synergy 3D map with positive ZIP indicating additive/synergistic effect and negative ZIP indicating inhibition, HN31 (E) ZIP mean= 2.7, p= 5.82e-01 and HN30 (F) mean= −2.23 (p= 7.62e-01).

**Supplemental Figure 2.**
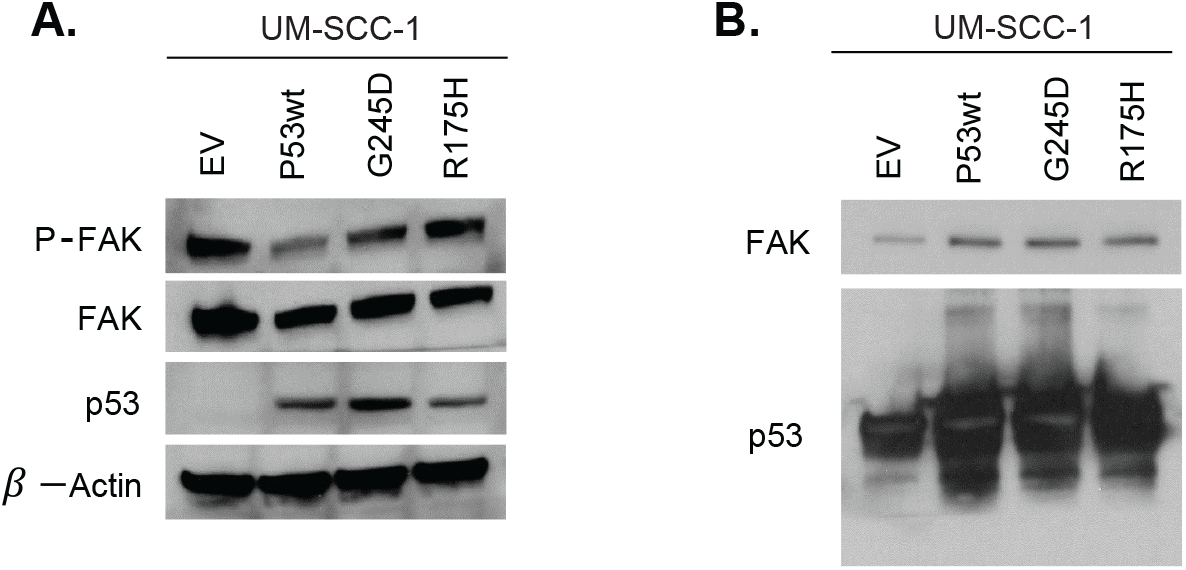
FAK activity and binding based on TP53 mutation. A) Immunoblot for the indicated proteins in UM-SCC-1 cells (p53 null/lo) expressing empty vector (EV), wild-type p53 (P53wt) or two missense TP53 mutations (G245D and R175H). B) Immunoprecipitation with p53 antibody in cell lines from (A) was performed as described in the Methods. The resulting immunoprecipitate was evaluated of total FAK and p53 via immunoblot.

